# Sexual signal reliability in male zebra finches: food intake explains the impact of immune activation on carotenoid-based coloration

**DOI:** 10.1101/2025.04.08.647553

**Authors:** Alejandro Cantarero, Rafael Mateo, Pablo Camarero, Pedro Andrade, Miguel Carneiro, Carlos Alonso-Alvarez

## Abstract

Theory predicts that traits evolving as animal signals must transmit reliable information about the bearer’s quality. Two hypotheses are often invoked to explain how carotenoid-based colorations function as reliable individual quality signals. The first hypothesis suggests that carotenoid allocation trade-offs between signaling and survival-related functions maintain signal reliability. Only high-quality individuals could resolve these trade-offs without incurring a fitness loss. Such signals are plastic, and the signaler can strategically manage their expression level to increase fitness returns. The second hypothesis posits that certain carotenoid-based signals are strongly constrained in their expression, not allowing strategic manipulations or cheating. We challenged captive male zebra finches (*Taeniopygia guttata*) with simulated infections by injecting lipopolysaccharides (LPS) derived from *Escherichia coli* and quantified food intake, carotenoid and vitamin levels in several tissues. Compared to controls, LPS-treated birds reduced the bill redness, a sexual signal. However, this reduction was largely explained by decreased food intake caused by LPS. Bill redness negatively correlated with body temperature and spleen mass but positively associated with blood, spleen and subcutaneous fat carotenoid levels. Overall, our findings suggest that carotenoid-based bill coloration predominantly reflects the recent physiological condition of individuals and resource allocation trade-offs rather than directly signaling individual quality.

## INTRODUCTION

Conspicuous traits in animals often function as signals in social and sexual selection contexts. For these traits to evolve as signals, they must reliably convey information that benefits both the signaler and the receiver (Laidre & Johnstone, 2013; Biernaskie et al., 2014). Typically, the information content of the signal reflects the individual quality of the signal bearer, thereby influencing the receiver’s mating and reproductive decisions. Among the traits commonly involved in sexual signaling are colorations, particularly those produced by pigments ranging from yellow to red. In vertebrates, many of these colorations are produced by carotenoids (e.g. Grether, 2004; McGraw, 2006). Carotenoids are pigments that must be obtained through food, and some authors suggested that they may be scarce in the environment, requiring specific foraging efforts to acquire them (Endler, 1980; Kodric-Brown, 1985; Hill, 1990). Moreover, carotenoids have important physiological functions as immune enhancers and antioxidants (Pérez-Rodríguez, 2009; Svensson & Wong, 2011). It has been suggested that carotenoid-based colorations are reliable individual quality signals as their production could incur costs (e.g., Weaver et al. 2017) or involve resource allocation trade-offs that require optimization (Szamado et al., 2022, 2023a). Both investing extra time and energy in foraging carotenoids, and the allocation of the ingested carotenoids to signals instead of homeostasis, may impose fitness costs or give place to suboptimal fitness returns (Møller et al., 2000; Alonso-Alvarez et al., 2008; but see Koch & Hill 2018). High-quality individuals are expected to better cope with these challenges, thereby expressing more intense coloration than low-quality individuals. In this scenario, theory suggests that the signal production mechanism should be enough plastic to allow animals to strategically make allocation decisions that optimize their fitness returns and potentially deceive signal receivers (e.g., Maynard Smith & Harper, 2003; Szamado et al., 2023b).

Nonetheless, according to animal signaling theory, some signals may be tightly constrained in their expression, exhibiting low plasticity and being strongly determined by individual quality (e.g. Maynard Smith and Harper, 2003). Such signals are theoretically unfalsifiable and are commonly referred to as “index signals” (Maynard Smith & Harper, 2003; Laidre & Johnstone, 2013; Biernaskie et al., 2014). It has been proposed that traits coloured by red ketocarotenoids (e.g. astaxanthin, α-doradexanthin, adonirubin, echinenone), which are enzymatically synthesized by some animals from dietary yellow carotenoids (xanthophylls such as lutein, zeaxanthin or β-cryptoxanthin), may serve as index signals (Hill, 2011; Lopes et al., 2016; Mundy et al., 2016; Weaver et al., 2017; Toomey et al., 2022). According to this hypothesis, the enzymes responsible for carotenoid conversion should be placed in the inner mitochondrial membrane and are tightly linked to cellular respiration processes (Johnson & Hill, 2013; Hill et al., 2019). This close association is thought to limit the potential for signal dishonesty, as the ability to produce red ketocarotenoids would inherently reflect the individual’s metabolic efficiency (Weaver et al., 2018; but see also Koch et al., 2025).

In birds, experimental immune activation, such as exposure to antigens, has been shown to reduce the expression of carotenoid-based signals (e.g., Fitze et al., 2007; Merrill et al., 2016; George et al., 2017). Among the most commonly used antigens in such studies are lipopolysaccharides (LPS) of the bacterial cell wall typically derived from gram-negative bacteria such as *Escherichia coli* or *Salmonella* spp. (e.g. Buras et al. 2005). LPS antigens trigger an innate immune response involving the cell and humoral immune system (Bryant et al., 2010; Rathinam et al., 2019; Scalf et al., 2019). In poultry, LPS exposure has been shown to induce inflammation and increase the mass of key immune and metabolic organs, particularly the liver (e.g., Xie et al. 2000; Liebolt et al., 2016; Bai et al., 2019) and spleen (e.g., Koutsos et al., 2003; Rajput et al., 2013; Liebolt et al., 2016; Bai et al., 2019; but see also Li et al., 2017). Moreover, experimental studies have consistently confirmed a negative impact of LPS in the carotenoid-based coloration and circulating carotenoid levels in avian species (Alonso-Alvarez et al., 2004; Torres & Velando, 2007; Gautier et al., 2008; Cote et al., 2010; Romano et al., 2011; Sild et al., 2011; Rosenthal et al., 2012; Csernus et al., 2020). Similar declines in carotenoid-based coloration have also been observed in reptiles (López et al., 2009; Ibañez et al., 2014) and even in humans (Henderson et al., 2017).

Here, we investigate the hypotheses concerning the reliability of carotenoid-based signals by analyzing carotenoid concentrations across multiple tissues (the bill, liver, subcutaneous fat, and spleen) of males of zebra finches (*Taeniopygia guttata*). This species possesses a sexually selected trait, the red bill, which is involved in mate choice (Simons & Verhulst, 2011) and is pigmented by endogenously converted ketocarotenoids (McGraw & Toomey, 2010). We used captive birds exposed to a simulated one-month infection via repeated LPS injections. Our experimental design was similar to that of a study conducted twenty years ago (the exact LPS dosage and very similar time-lapse; i.e., Alonso-Alvarez et al., 2004). In that study, birds were supplemented with carotenoids. Here, we decided to simplify the design, avoiding pigment supplementation as carotenoids are usually abundant in commercial diets (see here Supplementary Material; SM), and pigment addition has been shown to not proportionally increase signal expression (Koch et al., 2016). Alonso-Alvarez et al. (2004) reported significant declines in both bill coloration and blood carotenoid levels in LPS-treated zebra finches compared to controls. The same findings have been replicated two additional independent studies on this species (i.e., Gautier et al., 2008 and Cote et al., 2010). Here, however, we tested the influence of resource acquisition on signal expression by quantifying the ingestion of food (seed mix) containing yellow substrate carotenoids. We should note that LPS often inhibit food ingestion in different bird species (Baert et al., 2005; Rosenthal et al., 2012; Teemer & Hawley, 2024; but see Koch et al., 2018), including zebra finches (Burness et al., 2010; Sköld-Chiriac et al., 2015; Tapper et al., 2021). Therefore, the reduction in blood carotenoid levels and coloration previously described in LPS-treated birds (above) may be partly due to LPS-induced hypophagia, leading to decrease carotenoid intake. Here, we specifically address this point by evaluating the relationship between food consumption and the signaling outcomes.

We predict that individuals facing an immune challenge should prioritize self-maintenance by allocating carotenoids to vital organs, such as the liver to protect it against immune-induced damage (e.g., Kaulmann and Bohn 2014; Elvira-Torales et al., 2019; Abou Elazab et al., 2022) and spleen for favoring its immune activity (e.g., Milani et al., 2017; Csernus 2020, 2023), rather than investing them on coloration. Thus, mammalian studies suggest that inflammatory responses sequestrate retinol in the hepatic tissue (Rubin et al., 2017), and we may infer that something similar could occur with carotenoids as closely related compounds (Tang & Rusell, 2009). Alternatively, liver carotenoids might be consumed to counteract the oxidative stress linked to LPS-induced inflammation (e.g. Meriwether et al., 2010; Kaulmann & Bohn, 2014). Concerning the spleen, carotenoids have been shown to enhance spleen antibody production in mammals (reviewed, e.g., Milani et al., 2017) and to reduce LPS-induced spleen inflammation in chickens (Csernus 2020, 2023). Therefore, increasing spleen carotenoid levels should be an optimal strategy for combating an infection. For subcutaneous fat, we may predict that carotenoid levels may decrease in LPS-treated birds as a consequence of oxidative stress in adipose tissues (e.g., Bonet et al., 2015; Mounien & Tourniare, 2019). Finally, we hypothesize that food intake will be positively associated with signal intensity and carotenoid accumulation in ornaments and other tissues. However, the food intake and the immune challenge could also interact. In such a case, we predict that the relationship between food intake and signal intensity may be weaker in LPS-treated birds due to increased carotenoid demands for homeostasis.

Interpreting the results within the framework of the trade-off versus index signal hypotheses is challenging. We must first assume that individuals exhibiting the reddest bills at the start of the experiment represent high-quality individuals. Accordingly, low-quality individuals should invest more in self-maintenance and less in signal expression when enduring the immune challenge. In this case, initial redness should interact with treatment on the response variables. If such an interaction is not observed, it may suggest the occurrence of signal dishonesty or ‘cheating’, which would align more closely with the plastic resource allocation strategies proposed by the trade-off hypothesis.

### Material and Methods

Adult male zebra finches (*n* = 70) were individually housed in cages (0.6 m × 0.4 m × 0.4 m) placed in an indoor aviary (FIEB Foundation, Casarrubios del Monte, Spain) under controlled conditions (see SM). All birds received *ad libitum* water, grit, and a commercial seed mix (KIKI® exotic birds, Spain; composition in SM). After a minimum 14-day quarantine and acclimation period, body mass and bill redness were measured, and a blood sample was taken (Chronogram in Figure S1 of SM). Two days later, each feeder was replenished and then weighed. The feeders (ref. M051, STA Soluzioni, Italy) were equipped with a shell collector at the bottom, and an additional collector was placed at the entrance. As no seeds or shells were found on the floor of the aviary or cage, we are confident that all consumed grain was accurately quantified. The feeders were replenished and weighed every three days with the content of both collectors included in each measurement. The difference between two consecutive weights represented the amount of grain consumed during that period. The first three-day period served to test for a difference in food intake before the experiment began (i.e., before the first injection; below). The total grain mass consumed by each bird from the first injection day until two days before the last injection was considered the individual food intake over the 27-day experimental period. This value was corrected for individual body mass at the first injection day (i.e., “initial body mass”; Figure S2). Water expenditure (mL) was estimated by weighing water dispensers one day after the second injection and again three days later. Note that this measure may be partially influenced by water loss due to behaviors other than drinking.

Five days after blood sampling, all birds were assigned to treatments. Thirty-five birds received intraperitoneal injections of 0.01 mg of lipopolysaccharide (LPS) from the cell wall of *Escherichia coli* (serotype O55:B5; SIGMA, St. Louis) diluted in 0.1 mL of PBS (concentration = 0.1 mg/mL; i.e., LPS-treated birds). The remaining birds (n = 35) were injected with the same volume (0.1 mL) of PBS alone (control birds). The treatments were randomly distributed across aviary cages (rows and columns). No significant pre-treatment differences were found in body mass, body size (tarsus length), size-corrected body mass (body mass controlled for tarsus length, bill color measures (hue, saturation or brightness) or initial (pre-experimental) food intake (all *p*-values > 0.38). To maintain the immune challenge, birds were injected once per week for five consecutive weeks. Bill coloration and body mass measures were repeated 28 days after the first injection and the birds were assigned to one of three sampling blocks (*n* = 23, 23 and 24; treatment balanced among blocks: *χ*^2^ = 0.09, df = 2, *p* = 0.957). The blocks were necessary due to time limitations during tissue extraction. On day 28, birds in the first block were injected (fifth dose; Figure S1). The following day, a (second) blood sample was collected from these birds, after which the birds were euthanized via cervical dislocation. This procedure was repeated for the second and third blocks on subsequent days. Since the duration of the experiment differed among blocks (24-48h difference; Figure S1), the block was included as a random effect in the statistical models (see below).

Body temperature was measured immediately after euthanasia using an infrared thermometer. For the first nine birds, temperature was measured individuals at the midpoint of the breast after blowing feathers apart. However, these readings yielded unexpectedly low values (mean ± SD: 37.72 ± 1.46 °C, range: 34.7-39.9 °C). For the following birds, we directed the thermometer to pectoral muscles after being feather plucked (41.45 ± 0.94 °C; 39.6-42.9 °C). We standardized the values to obtain mean = 0 and SD = 1 in both groups. Std-values were then used in the statistical analyses. Similar results were obtained when analyses were restricted to the raw data from the latter group alone (*n* = 61; SM). Following euthanasia, tissues including the bill (ramphoteca, epidermis, and dermis), liver, spleen, intestines and subcutaneous fat (visible adipose tissue connected to the skin) were collected. An aliquot was obtained from the bill, liver, spleen, and fat for carotenoid and vitamin quantification using HPLC (see below). The samples were immediately snap-frozen in dry ice and stored at −80°C. The experiment was approved by the animal ethics government office (i.e., Consejería de Agricultura, Junta de Castilla La Mancha; approval ref. JCCM 27-2021) in agreement with Directive 2010/63/EU.

#### Color measurements

Digital photography (Nikon® D300) was used to measure bill redness. Each bird was photographed laterally, taking images from both sides of the head (see SM for details). Digital photographs were standardized and analyzed using ‘SpotEgg’ software (Gómez & Liñán-Cembrano, 2017). This image-processing tool for automatized analysis solves the need for linearizing the camera’s response to subtle light intensity variations (Stevens et al., 2007). Mean red, green, and blue (RGB) values measured from the lateral area of the bill (upper and lower mandibles) were used to calculate hue and brightness values following Foley & van Dam (1982). The exact measured area was recorded. Values from both sides of the bill were combined to obtain a single global measure. Measured repeatability (Lessells and Boag 1987), calculated on a subset (*n* = 30) of digital photographs measured twice was high (*r* > 0.90, *p* < 0.001). As lower hue values correspond to redder coloration, hue values were reversed by multiplying by -1. This transformed value was defined as “bill redness” and used in subsequent analyses.

#### Carotenoid, retinoid, and tocopherol analyses

Carotenoids, retinoids, and tocopherols were extracted following the protocols described by García-de Blas et al. (2013) and Rodríguez-Estival et al. (2010) (see SM for additional details on the carotenoid extraction). Identification and quantification were performed using a high-performance liquid chromatography system (HPLC) (Agilent Technologies 1200 series) equipped with diode array and fluorescence detectors and a C30 column (ProntoSIL 200-3-C30 3μm 250x4,6mm, Sharlab) (García-de Blas et al. 2011., 2013; Alonso-Alvarez et al., 2018). Standards of lutein, zeaxanthin, canthaxanthin, astaxanthin, 3HOE, β-cryptoxanthin, and β-carotene were obtained from CaroteNature (Lupsingen, Switzerland). Saponification with KOH did not yield additional carotenoid recovery (see saponification method in García-de Blas et al. 2011). Retinyl acetate was used as an internal standard (IS) for carotenoid and retinol quantification, whereas tocopheryl acetate was used as IS for tocopherol quantification. Concentrations were calculated with standard calibration curves (ranging from 0.5 to 15 nmol/tube) controlling for the peak area of the respective IS. Compounds were identified by comparing their retention times and UV-Vis spectra with those of standards and literature-reported spectra. Lutein derivatives detected in plasma samples were quantified using the lutein calibration curve, assuming identical response factors. Carotenoid concentrations were expressed in nmol/mL in the plasma and nmol/g in the liver, fat, spleen, and bill. Total carotenoid levels were calculated as the sum of all identified carotenoid concentrations within each sample. Blood carotenoid and vitamin levels (initial and final values) of the same bird were analyzed in the same laboratory session. Spleen carotenoid and vitamin levels could only be measured in a subset of samples (30%), corresponding to those that reached the required minimum mass (> 13 mg; *n* = 21; 10 controls and 11 LPS-treated birds). We should note that another aliquot of this tissue was reserved for RNA assays (unpublished material). The spleen carotenoid analyses were performed in a single laboratory session, with an even distribution of individuals across sampling blocks (*χ*^2^ = 0.04, df= 2, *p* = 0.979). As a result, these factors were not included as random effects in the statistical models.

#### Statistics

The values of bill redness (reversed hue), carotenoid and vitamin levels, organ mass and body temperature were tested as dependent variables in separate models to disentangle resource allocation strategies. ANCOVAs were used for bill redness, organ mass, and body temperature, while Generalized Linear Mixed Models [GLMM] were applied to most molecule concentration variables (i.e. PROC MIXED in SAS version 29). The laboratory session and the sacrifice block were added as random terms in the GLMMs; however, their influence was consistently non-significant (*p* > 0.08 for all tests). For fat and spleen compounds, no influence of the random terms was detected (*p* > 0.90), and we used Generalized Linear Models (GLMs).

Dependent variables were log- or square-root transformed when necessary to meet normality assumptions. When transformed data still violated normality criteria non-parametric statistical tests were applied. The distribution of subcutaneous fat retinol and carotenoid levels was zero-inflated (i.e., 24% and 34% zero values, respectively). To account for this, we applied zero-inflated negative binomial (ZINB) generalized linear models (GLMs) (PROC GENMOD in SAS v29). The ZINB GLMs separately analyze the normal and zero-inflated parts of the distribution (Ridout et al., 1998; Cantarero et al., 2020).

In all models, we followed a stepwise approach. First, we tested the effect of the treatment alone. In the second step, we included treatment, initial bill redness (covariate), and their interaction. In the third step, we tested treatment, food intake (as a covariate), and its interaction with treatment. The “initial bill redness” covariate was the standardized residual of a linear regression testing the initial bill hue (reversed) on two confounding covariates: initial bill brightness (slope ± SD: = -8.28 ± 0.96, *t* = 8.59, *p* < 0.001) and the bill area selected for the measurement (slope ± SD: 1.58 ± 0.499, *t* = 3.15, *p* = 0.002). The “food intake” covariate was the standardized residual of a linear regression testing total food intake on initial body mass (slope ± SD: 2.29 ± 0.55, *R^2^* = 0.20, *p* < 0.001; Figure S2 in SM). The results remained qualitatively unchanged if food intake values were uncorrected by body mass when tested as a covariate. The “final bill redness” term was used to define reversed hue corrected for bill hue and bill measured area at the last sampling, i.e., not corrected for initial variability.

To account for potential confounding effects, additional covariates were included in specific models. For the bill redness model, we included bill brightness and bill area (both measured at the final sampling) along with the initial bill redness covariate. The model testing plasma carotenoids included the “initial carotenoid values” that were, in turn, the standardized residuals from a GLMM testing initial values as the dependent variable and the laboratory session and the block of sacrifice as random factors.

The correlation coefficients for the association between bill redness and total carotenoid levels in each tissue were calculated by testing the standardized residuals of each parameter obtained from GLMMs (i.e., excluding the treatment but maintaining confounding covariates and random factors). For the spleen, where measurements were available only for a reduced sample (n = 21), simpler statistical tests were applied depending on data distribution (Student’s t-test, ANCOVA, or Mann–Whitney U tests).

Two-tailed statistics were used for most *p*-value calculations as the direction of the effect was not well-established by previous studies. However, one-tailed *p*-values were reported when assessing the effect of LPS treatment (not its interaction) on bill color and plasma total carotenoid levels, given the substantial prior evidence supporting a clear directional prediction (i.e., an LPS-induced decrease in their values; see Introduction; see also Rice and Gaines 1994; Ruxton & Neuhaüser, 2010; Cho & Abe, 2013). Satterthwaite’s correction was used to calculate degrees of freedom in GLMMs. Effect sizes are reported as Cohen’s *d* for group comparisons with *p*-values < 0.12, while Pearson’s correlation coefficients are used to represent effect sizes in correlational analyses (Cohen, 1992 In instances where carotenoid levels could not be quantified due to extraction issues (see sample sizes in Table S1), the covariates (food intake or initial bill redness residuals) were recalculated from the dataset that only included values for the dependent variable.

## RESULTS

### Water, food intake and body mass

The LPS-treated birds exhibited significantly higher (log-transformed) water expenditure than controls (Student’s *t*: 2.01, df = 54.7, *p* = 0.049, *n* = 69; Figure 1A for raw values). LPS-treated birds also showed lower food intake (*F*_1,67_ = 4.49, *p* = 0.038, *n* = 70; initial body mass: *F*_1,67_ = 18,01, *p* < 0.001, estimate ± SE: 2.29 ± 0.540). The immune challenge did not influence final body mass (treatment: *F*_1,67_ = 0.28, *p* = 0.597; initial body mass: *F*_1,67_ = 1088, *p* < 0.001; estimate ± SE: 1.04 ± 0.032). Including food intake as an additional covariate did not significantly explain final body mass variability (*p* = 0.434). Neither the initial bill redness (*p* = 0.433) nor its interaction with treatment (*p* = 0.802) significantly influenced food intake.

**Figure 1.**
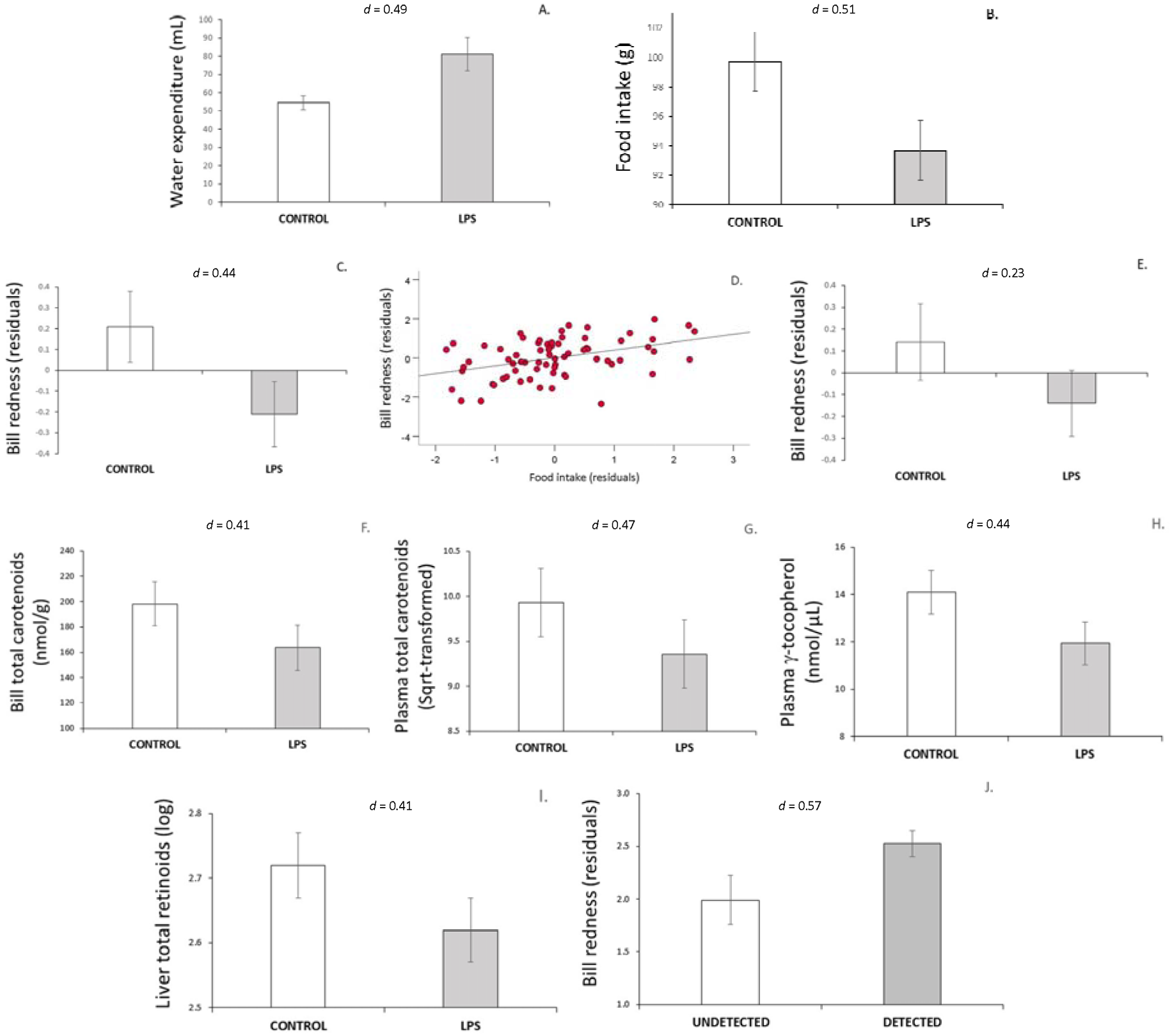
LPS impact on water expenditure (A), food intake (B) and bill redness (C; i.e., residuals of a mixed model controlling for bill brightness, bill measured area, initial bill redness and sacrifice session). Relationship between bill redness (those residuals) and food intake controlled for initial body mass (residuals) (D). LPS impact on bill redness after controlling for food intake residuals (E), bill total carotenoid levels (F), plasma total carotenoid (G), plasma γ-tocopherol (H) and liver total retinoid levels (I). Difference in bill redness (Table 1 model’s residuals, see Results) in birds with undetectable or detectable carotenoids in their subcutaneous fat (L). Least squared means ± SE from mixed models (other comparisons: means ± SE). Cohen’s *d* effect sizes are shown.

### Bill color

At the end of the experiment, LPS-treated birds exhibited lower bill redness compared to controls (treatment: *F*_1,65_ = 3.21, *p* = 0.039; brightness: *F*_1,65_ = 28.4, *p* < 0.001, estimate ± SE: -4.45 ± 0.835; area measured: *F*_1,65_ = 15.29, *p* < 0.001, estimate ± SE: 1.86 ± 0.475; initial redness: *F*_1,65_ = 13.81, *p* < 0.001, estimate ± SE: -1.043 ± 0.281; see Figure 1C). The interaction between initial bill redness and treatment was non-significant when added to the same model (Table 1). However, when the food intake covariate was also added, it showed a significant positive correlation to bill redness (Figure 1D) without interaction with treatment (Table 1), whereas the latter became non-significant (Table 1; Figure 1E). The correlation between the treatment and food intake remained significant when the interaction was removed (i.e., *F*_1,64_ = 11.35, *p* = 0.001; estimate ± SE: 0.923 ± 0.274).

**Table 1.**
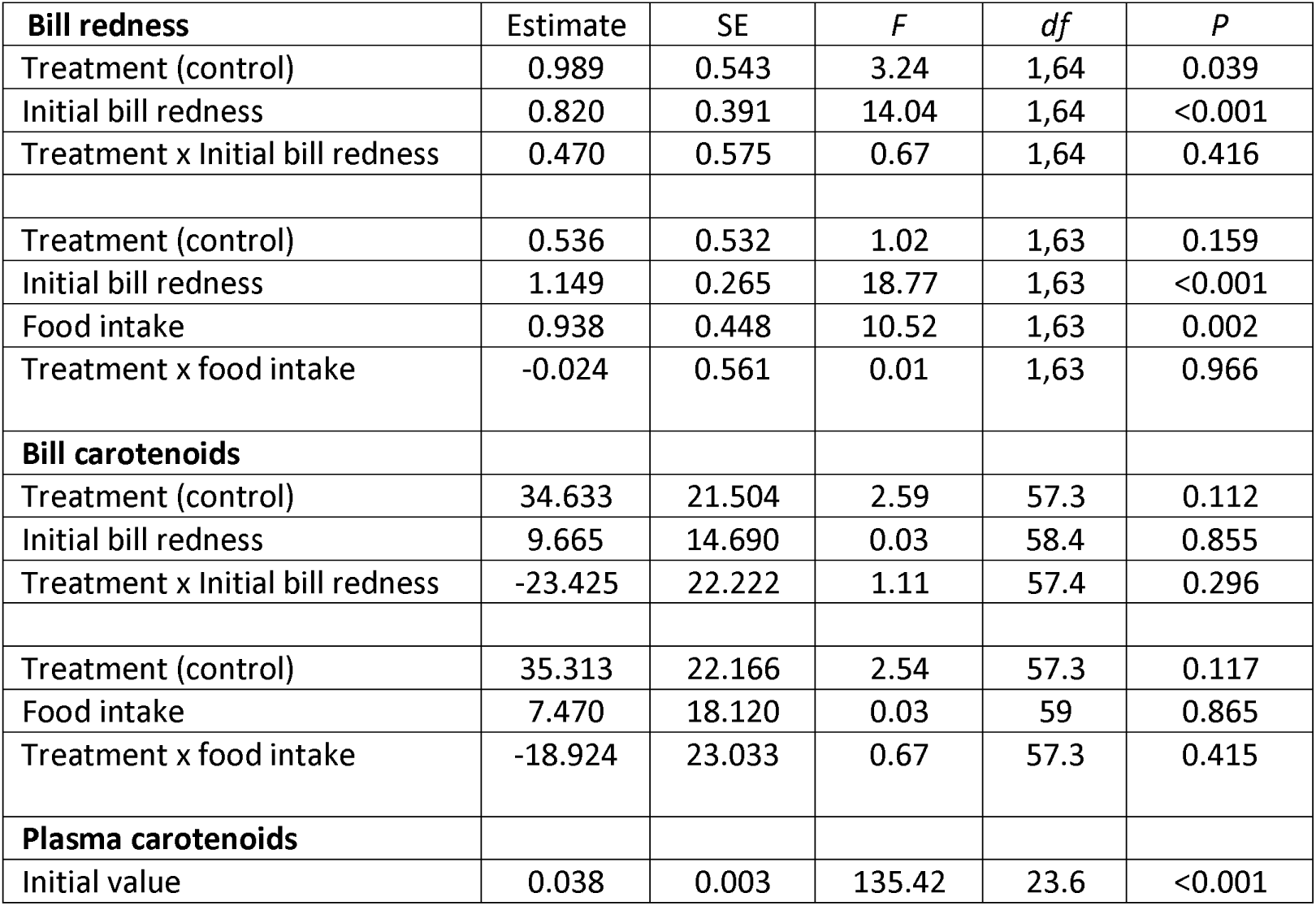
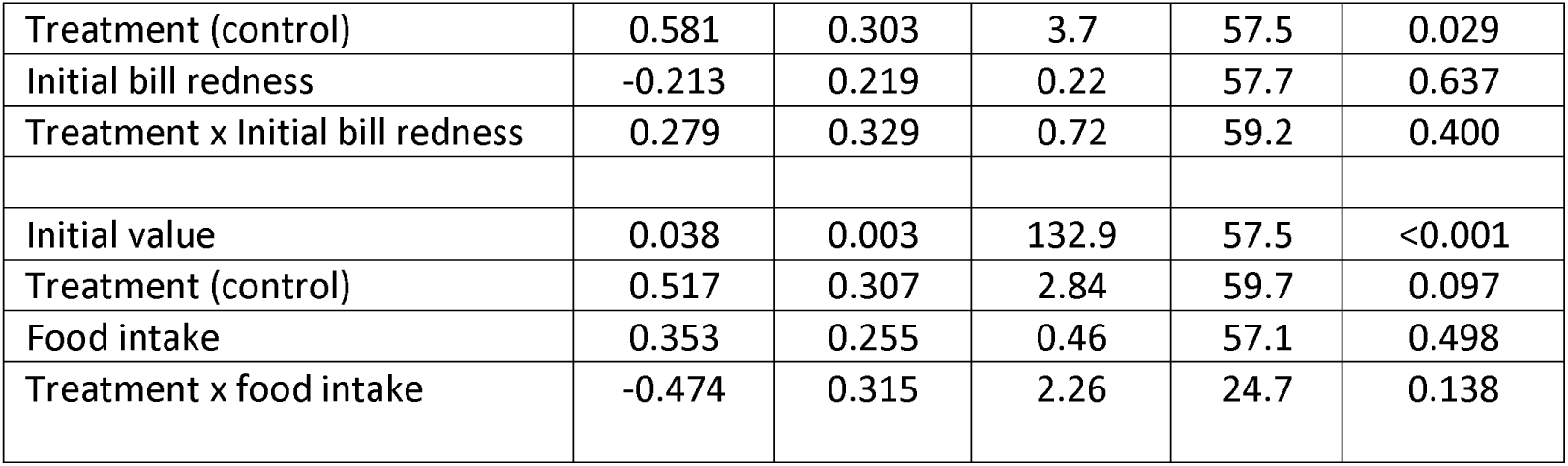
Models testing the impact of the immune challenge on bill redness (*n* = 70), and bill (*n* = 63) and plasma (*n* = 67) carotenoid levels. The bill redness models included bill brightness and measured area as covariates (both p < 0.001). The initial bill redness covariate was the residual of a model testing bill redness at the start of the experiment corrected by bill brightness and measured area at the same time. One-tailed *p*-values for the treatment factor on bill redness and plasma carotenoids (see Statistical analyses).

### Bill composition

In the bill, we detected retinol, astaxanthin, α-doradexanthin, adonixanthin, zeaxanthin, α- and γ- tocopherols, as well as two unidentified carotenoids (Table S1). LPS-treated birds exhibited lower mean total carotenoid levels than controls (Figure 1F); however, this treatment effect only showed a weak trend toward significance (*F*_1,59.3_ = 2.63, *p* = 0.110). The inclusion of the initial bill redness or food intake covariates and their interactions with treatment did not yield significant effects (Table 1). Retinol was detected in nine birds, evenly distributed across treatments (*χ*^2^ = 2.13, df= 1, *p* = 0.144), with no significant difference between groups (Mann-Whitney *U* = 519, *p* = 0.219). The two tocopherols did not differ between treatments (*F*’s *p*-values > 0.18). Bill total carotenoid levels (residuals from the Table 1 model without treatment) were positively correlated to bill redness when assessed as reversed hue at the end of the experiment (Pearson’s *r* = 0.27, *p* = 0.033, *n* = 63; Figure S3).

### Blood plasma

In plasma samples, we detected retinol, lutein, zeaxanthin, four lutein derivatives, echinenone (in only two samples), and α- and γ–tocopherols (Table S1). LPS-treated birds showed significantly lower total carotenoid levels (square-root transformed) than controls (treatment: *F*_59.4_= 3.67, *p* = 0.030; initial value: *F*_1,23.5_ = 140.12, *p* < 0.001, estimate ± SE: -0.038 ± 0.003; Figure 1G). Neither initial bill redness, food intake, nor their interaction with treatment significantly explain variation in plasma total carotenoid levels (Table 1).

Given the rapid turnover of circulating carotenoids, we further tested food intake recorded exclusively during the week prior to the last blood sampling (mean ± SD: 21.14 ± 3.02 g). In this case, a significant interaction with treatment emerged (treatment: *F*_1.57.2_ = 7.73, *p* = 0.004; initial value: *F*_1,25.5_ = 135.28, *p* < 0.001; food intake: *F*_1.57.7_ = 0.54, *p* = 0.465; interaction: *F*_1.57.3_ = 6.48, *p* = 0.014). Specifically, a positive relationship between food intake and plasma carotenoid levels was found in LPS-treated birds (slope ± SE: 0.164 ± 0.055; *F*_1.24.9_ = 8.77, *p* = 0.007; *r* = 0.49) but not in controls (−0.105 ± 0.085; *F*_1.29.6_ = 1.54, *p* = 0.225; *r* = -0.09).

When each carotenoid was tested individually, all showed at least a statistical trend towards reduced levels in LPS-treated birds (all *p* < 0.10), except zeaxanthin (i.e., *p* = 0.675). Furthermore, total plasma carotenoid levels at the last blood sampling (residuals from a mixed model only controlling for random terms) were positively correlated with final bill redness (Pearson’s *r* = 0.33, *p* = 0.006, *n* = 70). Retinol or α-tocopherol levels did not differ significantly between treatments (*p*-values > 0.66). However, LPS-treated birds showed a statistical trend toward lower plasma γ-tocopherol levels compared to controls (*F*_1.61.9_= 3.22. *p* = 0.077; initial value: *F*_1.28.1_ = 1975. *p* < 0.001; estimate ± SE: 0.986 ± 0.022; Figure 1H). The addition of the food intake covariate to this model did not report a significant effect (*p* = 0.461).

### Liver

The relative liver mass (i.e., controlled for body mass) did not differ significantly between treatments (ANCOVA: *F*_1.67 =_ 1.85, *p* = 0.178, *n* = 70; body mass at sacrifice: *F*_1.67 =_ 27.57, *p* < 0.001). The initial bill redness and its interaction with treatment had no significant effect on relative liver mass (all *p*-values > 0.60). Similarly, treatment had no significant effect on total carotenoid levels (*F*_1.63 =_ 0.01, *p* = 0.996). Neither the initial bill redness nor its interaction with treatment influenced liver total carotenoid levels (Table 2). The interaction between food intake and treatment was also non-significant; however, food intake alone showed a trend toward a positive association with liver carotenoid levels (Table 2), suggesting that birds consuming more food tended to accumulate higher carotenoid levels in the liver (Figure S4).

**Table 2.** Models testing the impact of the immune challenge on the liver, subcutaneous fat, and spleen total carotenoid levels. Alternative models testing the influence of initial bill redness and food intake are shown.

| **Liver (Normal)** | **Estimate** | **SE** | **F** | **df** | **P** |
| --- | --- | --- | --- | --- | --- |
| Treatment (control) | -0.002 | 0.282 | 0.00 | 61 | 0.993 |
| Initial bill redness | -0.121 | 0.209 | 0.19 | 61.1 | 0.669 |
| Treatment x Initial bill redness | 0.118 | 0.290 | 0.17 | 61.1 | 0.684 |
| Treatment (control) | -0.153 | 0.285 | 0.29 | 61.2 | 0.593 |
| Food intake | 0.388 | 0.250 | 3.85 | 61.7 | 0.054 |
| Treatment x food intake | -0.166 | 0.303 | 0.30 | 61.1 | 0.586 |
| **Subcutaneous fat (ZINB)** | **Estimate** | **SE** | **$\chi^2$** | **df** | **P** |
| Treatment (control) | -0.217 | 0.212 | 1.04 | 1 | 0.307 |
| Initial bill redness | 0.105 | 0.128 | 0.25 | 1 | 0.616 |
| Treatment x Initial bill redness | -0.111 | 0.199 | 0.32 | 1 | 0.573 |
| Treatment (control) | -0.221 | 0.210 | 1.11 | 1 | 0.292 |
| Food intake | 0.328 | 0.163 | 0.49 | 1 | 0.482 |
| Treatment x food intake | -0.488 | 0.238 | 4.22 | 1 | 0.040 |
| **Spleen (Normal)** | **Estimate** | **SE** | **F** | **df** | **P** |
| Treatment (control) | -2.850 | 4.578 | 0.388 | 1.17 | 0.542 |
| Initial bill redness | 1.664 | 3.974 | 0.004 | 1.17 | 0.948 |
| Treatment x Initial bill redness | 2.995 | 5.068 | 0.349 | 1.17 | 0.562 |
| Treatment (control) | -3.954 | 4.330 | 0.834 | 1.17 | 0.374 |
| Food intake | 4.913 | 4.034 | 1.563 | 1.17 | 0.228 |
| Treatment x food intake | -3.727 | 4.878 | 0.584 | 1.17 | 0.455 |

In terms of specific compounds, lutein, three lutein derivatives, zeaxanthin, and β-cryptoxanthin were detected in the liver. None of them showed significant differences between treatments when analyzed separately (all *p’s* > 0.30). Retinol, four unidentified retinoids, α- and γ-tocopherol were also detected (Table S1). Total retinoid levels (log-transformed) showed a weak trend toward a decrease in LPS-treated birds (*F*_1.62.5_ =2.76, *p* = 0.101; Figure 1-I). α- and γ-tocopherol did not differ significantly between treatment (*p*-values > 0.59). Finally, liver total carotenoids or retinoids (i.e., residuals from their mixed models without treatment) were not significantly correlated with bill redness (i.e., Table 1 model without treatment residuals; *r* = 0.132, *p* = 0.286, and *r =* -0.170, *p* = 0.169, respectively).

### Subcutaneous fat

We detected retinol, zeaxanthin, lutein, a lutein derivative, and α- and γ- tocopherols in fat(Table S1). Total carotenoid levels were not significantly affected by treatment (*χ*^2^ = 1.07, *p* = 0.301; zero-inflated part of the model: *χ*^2^ = 0.18, *p* = 0.675). Similarly, the initial bill redness and its interaction with treatment had no significant effect (Table 2). However, food intake showed a significant interaction with treatment in the continuous part of the model (Table 2). When analyzed separately, no significant association between food intake and fat carotenoids was found in controls (slope ± SE: -0.148 ± 0.141; *χ*^2^ = 1.10, *p* = 0.294), but LPS-treated birds exhibited a trend toward a significant positive relationship (slope ± SE: 0.331 ± 0.182; *χ*^2^ = 3.32, *p* = 0.068). The zero-inflated part of these models consistently reported non-significant results (*p* > 0.46). The analyses of each parameter separately did not show a significant treatment effect (all *p-values* > 0.20). Additionally, we tested whether the presence or absence of fat carotenoids (as a binary factor) predicted bill redness in the model from Table 1. Birds with detectable fat carotenoid levels showed significantly redder bills (*F*_1.64 =_ 5.76, *p* = 0.019; *d* = 0.57; Figure 1-J). The inclusion of this binary factor did not alter the treatment effect on bill color (*p* = 0.027) and showed no significant interaction with treatment (*p* = 0.980).

### Spleen

LPS-treated birds were characterized by significantly heavier spleens (i.e., “relative mass”: log-transformed spleen mass controlled for body mass) compared to controls (ANCOVA: *F*_1.67 =_ 4.82, *p* = 0.032, *n* = 70; body mass at sacrifice: *F*_1.67 =_ 11.04, *p* = 0.001, *d* = 0.52; least squared means ± SE: 1.01 ± 0.06 and 1.19 ± 0.06, respectively; Figure 2A for raw values). Initial bill redness was negatively associated with relative spleen mass (*F*_1.65 =_ 7.41, *p* = 0.008, slope ± SD: -0.162 ± 0.059), with no significant interaction with treatment (*F*_1.65 =_ 1.12, *p* = 0.293). The negative correlation persisted after removing the interaction and treatment factor (*F*_1.67 =_ 7.30, *p* = 0.009; *r* = -0.31, *n* = 70), indicating that birds with redder bills at the start of the study exhibited lighter spleens than paler individuals (Figure 2B). Food intake did not influence spleen mass nor interact with treatment (both *p-values* > 0.55). Additionally, relative spleen mass (residuals) had no significant effect when added as a covariate to the Table 1 bill redness model (*F*_1.64 =_ 2.10, *p* = 0.152).

**Figure 2.**
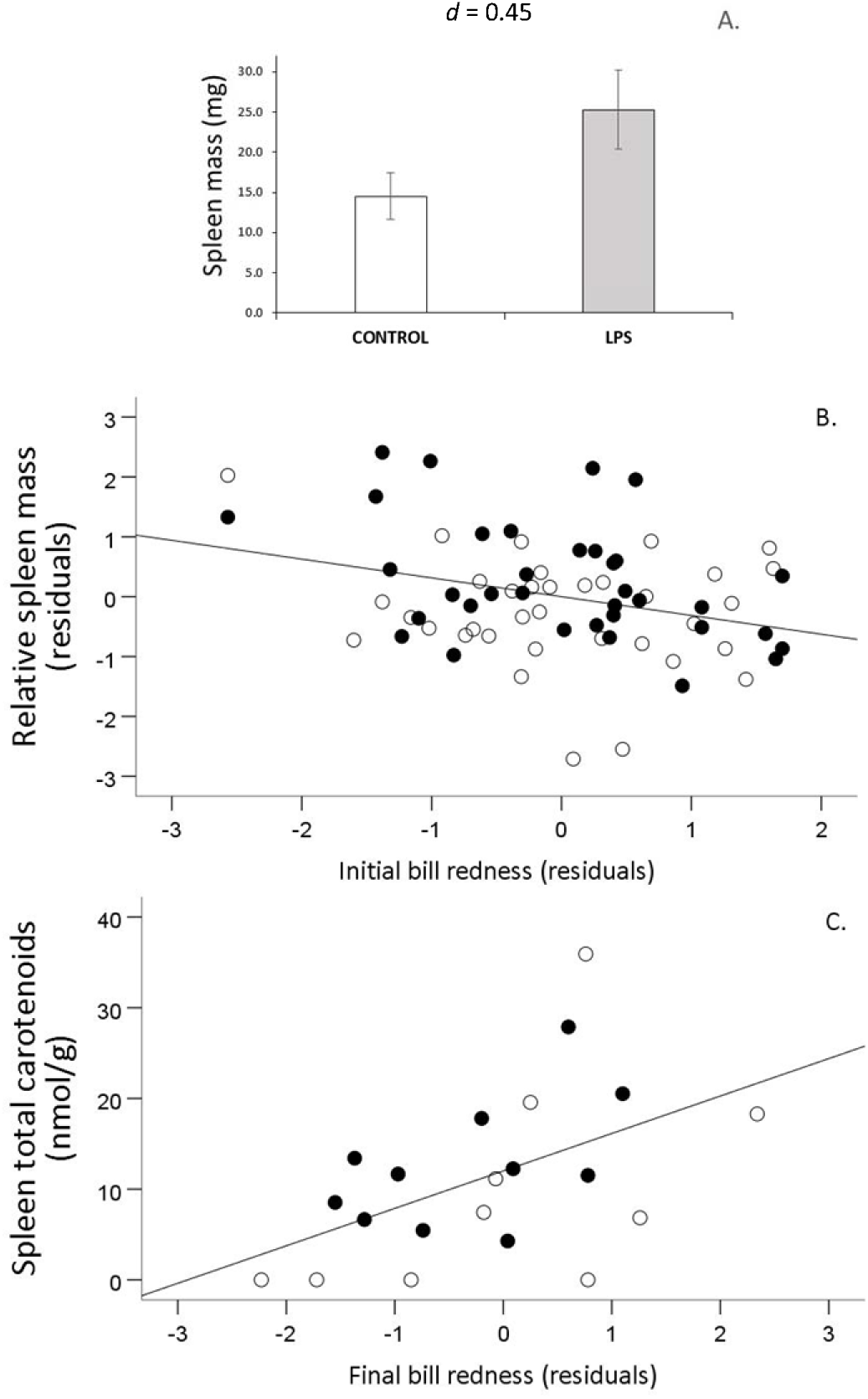
(A) Treatment effect on spleen mass (raw values; means ± SE). (B) Relationship between spleen mass controlled for body mass (residuals) and initial bill redness and (C) between spleen total carotenoid levels and final bill redness. Bill redness were residuals from models with bill brightness and measured bill area at the first (B) and second (C) sampling. Empty and filled dots for controls and LPS-treated birds, respectively. The treatment did not interact with bill redness (see Results).

Carotenoids in spleen tissue could only be analyzed in 10 controls and 11 LPS-treated birds that provided enough tissue mass (> 13 mg). These birds were significantly heavier at both the start and end of the experiment compared to the remaining individuals (Student’s *t* both: *p* < 0.010, *d* = 0.70 and 0.71, respectively). They also showed a trend to significantly lower initial bill redness compared to non-analyzed birds (*F*_1.66 =_ 3.65, *p* = 0.060, *d* = 0.47; means ±SE: -0.32 ± 0.25 and 0.14 ± 0.13, respectively). Note that birds with heavier spleens were associated with paler initial bill coloration in the whole of the sample (Figure 2B), a relationship also present in the subsample (*r* = -0.46, *P* = 0.035; *n* = 21; Figure S5).

Within this subsample, LPS-treated birds had higher average spleen masses than controls (mean, median, and rank: 43, 33, 96 mg and 30, 21, 86 mg, respectively), although the difference was non-significant (Mann-Whitnney’s *U* = 36, *p* = 0.181). The treatment did not affect initial bill redness (*F*_1.17_ = 1.54, *p* = 0.232) or bill redness controlled for initial color (*F*_1.16_ = 0.01, *p* = 0.936). β-cryptoxanthin, zeaxanthin, lutein, three lutein derivatives, retinol, three unidentified retinoids, and α- and γ-tocopherol were detected in spleen tissue (Table S1). Total carotenoid levels were unaffected by treatment (*t* = 0.67, df = 19, *p* = 0.512), initial bill redness, or food intake (Table 2). When testing individual compounds, only lutein showed a trend towards a treatment effect (*t* = 1.78, df = 19, *p* = 0.091, *d* = 0.77), with LPS-treated birds showing higher lutein values (means ± SE: 2.06 ± 1.90 and 1.21 ± 0.46 nmol/g, respectively). Interestingly, lutein was detected in all LPS-treated birds, whereas five control birds showed zero values (*χ*^2^ = 7.22, df=1, *p* = 0.007). A similar pattern was observed for total carotenoid levels, with four control birds lacking detectable carotenoids, compared to none in the LPS-treated group (*χ*^2^ = 5.44, df =1, *p* = 0.020).

Lastly, we tested whether spleen carotenoid content could be revealed by the concurrent bill redness measure. In a model analyzing spleen total carotenoids (dependent variable), final bill redness showed a significant positive association (*F*_1.17_ = 5.88, *p* = 0.027; slope ± SE: 4.12 ± 2.93; *r* = 0.50; Figure 2C) but no interaction with treatment (treatment: *F*_1.17_ = 1.23, *p* = 0.283; interaction: *F*_1.17_ = 0.02, *p* = 0.895). This positive relationship remained significant (i.e., *F*_1.16_ = 7.29, *p* = 0.016, slope ± SE: 4.27 ± 1.51) when adding the spleen mass as a covariate (*F*_1.16_ = 3.29, *p* = 0.089, slope ± SE: 0.12 ± 0.06) to test the absolute amount of spleen carotenoids.

### Body temperature

The body temperature was significantly higher in LPS-treated birds compared to controls (*t* = 2.18, df = 68, *p* = 0.032; Figure 3A). When the initial bill redness was included as a covariate in an ANCOVA model testing body temperature, a trend toward a significant interaction with treatment was observed (*F*_1.66_ = 0.03, *p* = 0.861; interaction: *F*_1.66_ = 3.47, *p* = 0.067). The sign of the slope differed between LPS-treated and control birds (slope ± SE: 0.197 ± 0.158 and -0.238 ± 0.172, respectively), but neither slope significantly deviated from zero (*p* > 0.175). When we instead tested the final bill redness, this covariate was negatively correlated (*F*_1.66_ = 4.37, *p* = 0.041; slope ± SE: -0.151 ± 0.178; *r* = 0.32; Figure 3B; interaction: *F*_1.66_ = 0.66, *p* = 0.419). Food intake did not affect body temperature nor interact with treatment (both *p-values* > 0.25). No significant associations between body temperature and carotenoid levels in the bill, plasma, spleen or fat were detected (all *p*-values > 0.22). Body temperature was also unrelated to relative spleen mass (*r* = 0.03, *p* = 0.820, *n* = 70).

**Figure 3.**
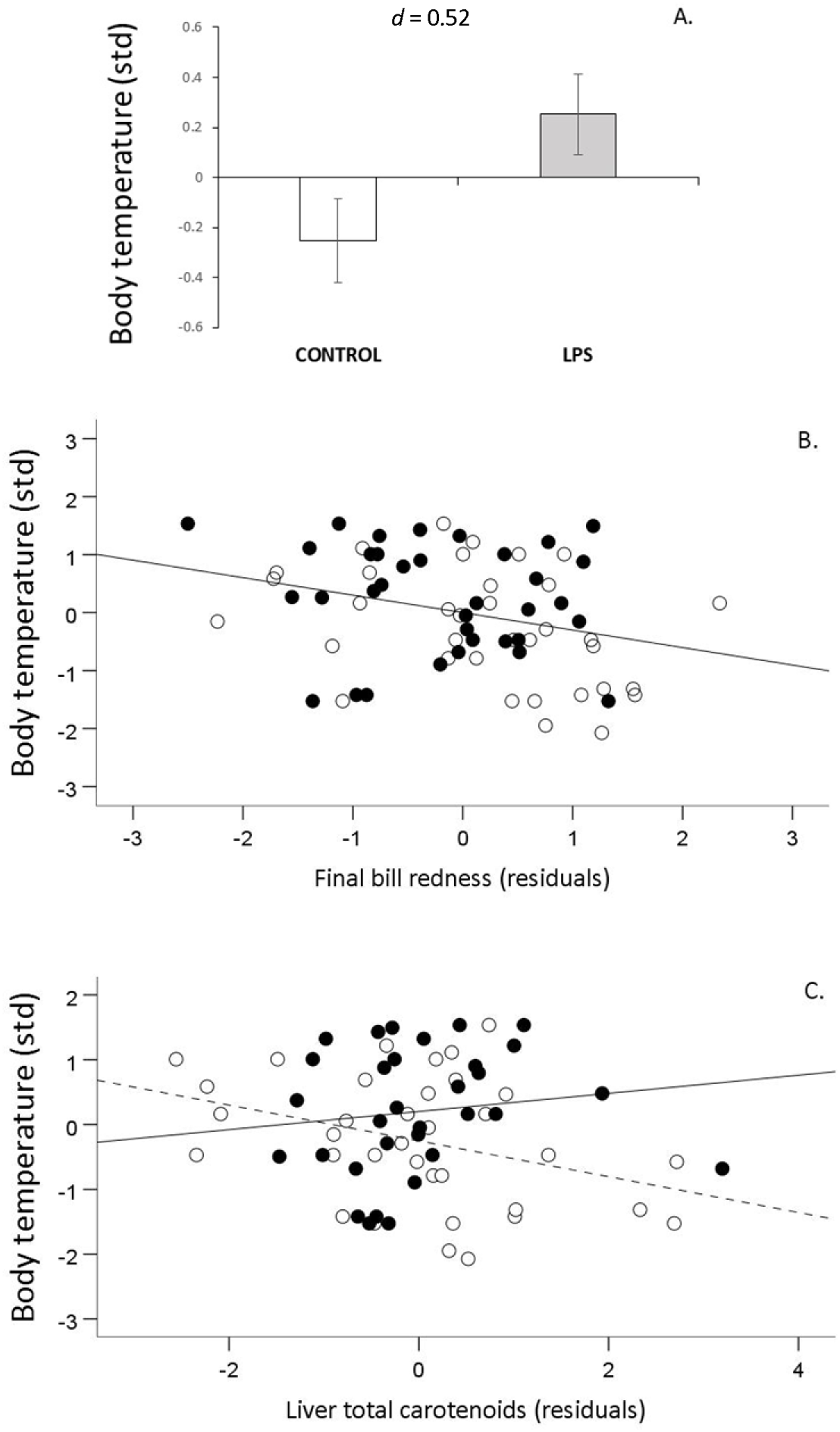
LPS effect on body temperature (A; means ± SE), relationship between body temperature and final bill redness (B) or liver total carotenoids (C). Temperature are standardized values and final bill redness were residuals from a mixed model including bill brightness and measured area at the last sampling event (see Methods). Controls: empty dots and dotted line. LPS-treated birds: filled dots and continuous line.

Nonetheless, when testing liver total carotenoid levels (residuals from a mixed model with only the random terms), we found a trend toward a significant interaction between body temperature and liver carotenoids (covariate: *F*_1.63_ = 0.37, *p* = 0.545; interaction: *F*_1.63_ = 3.46, *p* = 0.067). Control birds showed a negative relationship (*F*_1.33_ = 4.69, *p* = 0.038; slope ± SE: -0.276 ± 0.128; *r* = -0.35), but LPS-treated birds did not (*F*_1.30_ = 0.57, *p* = 0.457; slope ± SE: 0.140 ± 0.186, *r* = 0.14; Figure 3C).

## DISCUSSION

Our experimental immune challenge decreased bill redness and circulating carotenoid levels in male zebra finches, agreeing with results from previously published data in this and other species (see Introduction). However, we detected no significant interaction between initial bill redness and treatment effects, suggesting that the birds’ response to the immune challenge was not influenced by their initial signal expression. This implies that the strategy followed by the birds when investing in signaling under pressure (immune challenge) is independent of their previous condition. Thus, the link between signal level and individual quality could be weaker than expected for an index signal. Moreover, the amount of food intake strongly influenced the signal expression level, highlighting the high plasticity of the ketocarotenoid-based ornament.

The high plasticity of the zebra finch red bill can also be deduced from four experimental studies in this species reporting quick changes in the trait. Thus, an increase in bill redness was found in birds exposed to handling stress during four weeks (McGraw et al., 2011). A similar bill color increment was found in males housed with females during three weeks compared to those housed with males (Gautier et al., 2008). Even quicker, testosterone implants increased male bill redness in three days (Ardia et al., 2010) and certain immune activation (complete Freund’s adjuvant injection) declined bill color in only 48 hours (George et al., 2017).

Four findings support the presence of systemic inflammation in our LPS-treated birds. First, there was a trend towards a significant decrease in blood γ-tocopherol values. This could be attributed to its consumption when fighting the reactive nitrogen species (e.g., nitric oxide) released by inflammation (e.g., Christen et al., 1997; Jiang et al., 2022). Second, hepatic retinoids also showed a tendency to decline, which could also be due to their use as antioxidants (Tesoriere et al., 1993; Palace et al., 1999) or as a substrate for retinoic acid production, essential to mount the cellular immune response (Cifelli & Rose, 2006; Broadhurst et al., 2012; Rubin et al., 2017). Third, LPS-treated birds exhibited heavier spleens, consistent with previous results in poultry (Koutsos et al., 2003; Rajput et al., 2013; Bai et al., 2019). Fourth, LPS-treated birds showed higher body temperatures than controls, in agreement with findings in different passerine species, including zebra finches (Sköld-Chiriac et al., 2015; Tapper et al., 2021; but see Burness et al., 2010).

The LPS-induced reduction in food intake seems to have limited carotenoid allocation to sexual coloration. The potential inhibitory effect of LPS on food intake is currently well-known (see Introduction). We should highlight that an LPS-induced decline in carotenoid-based colorations was interpreted in this and other animal species as directly due to carotenoid being allocated to immune function stimulation or combat reactive oxygen species released by inflammation (e.g., Alonso-Alvarez et al., 2004; Torres & Velando, 2007; López et al., 2009; Cote et al., 2010; Romano et al., 2011; Rosenthal et al., 2012). However, our result highlights the necessity of controlling birds’ behavior when testing the impact of immune challenges on carotenoid-based ornaments. The possibility of a food intake influence on carotenoid-based signaling under an immune challenge or stress was previously argued (McGraw et al., 2011; Rosenthal et al., 2012) but has not been directly demonstrated until now.

The observed influence of food intake on bill redness challenges the notion that red ketocarotenoid-based signals depend on intricate production mechanisms (involving cell respiration) that are supposedly independent of resource allocation strategies (Johnson & Hill, 2013; Hill et al., 2019, 2023). While it is theoretically expected that such complex mechanisms ensure signal reliability through constraints (Podos, 2022), our findings suggest that resource acquisition can deeply influence carotenoid-based colorations, a possibility early suggested (Endler, 1980; Kodric-Brown, 1985; Hill, 1990; Møller et al., 2000). Specifically, male zebra finches with redder bills appear to be those better able to acquire sufficient carotenoid-rich food.

However, food intake itself was not directly correlated with bill carotenoid levels. Thus, the link between food intake and bill redness (Figure 1D) may not be uniquely attributed to dietary carotenoids acquired from the food. Perhaps, the overall nutritional state could have caused changes in the tissue microstructure affecting coloration (e.g., Siefferman & Hill 2005; D’Alba et al., 2014). Additionally, limitations in carotenoid extraction from the bill, which is a highly keratinized tissue, may contribute to the absence of significant correlations. For instance, in a similar trait (i.e., the bill of red-legged partridges, *Alectoris rufa*), we found a carotenoid recovery rate of 87% ± 3% (mean ± CV; García de Blas et al., 2013). This means that 10-15% of the bill pigments could not be extracted, adding error to the measure and complicating the detection of real effects.

Several findings support the reliability of red bill coloration as an indicator of individual condition in our study. First, intriguingly, the birds with the reddest bills at the start of the experiment had smaller spleens when sacrificed one month later, regardless of treatment (Figure 2A). A larger spleen is often interpreted as a sign of better immune competence (e.g., in birds: Møller & Erritzøe, 2002; Ardia, 2005; in mammals: Corbin et al., 2008). In that scenario, the red keto-carotenoid-based signal could cheat those interpreting it as redder bill individuals would have smaller spleens and, hence, poorer immune competence. Alternatively, the reddest bill birds could be the healthiest individuals if we consider that the spleen may become augmented during active infections (e.g., Smith & Hunt, 2004; Figuerola et al., 2005). Our birds previously passed a quarantine, and their behavior did not suggest health problems. In any case, we cannot discard the presence of subtle infectious processes.

More intuitive support for red bill signal reliability was that the birds with the reddest bills also had the lowest body temperatures. Here, we assume that the higher body temperatures in the reported range (e.g., > 42°C) indicate underlying inflammatory processes (raw values in Figure S7B, C). Body temperatures were also negatively correlated to liver total carotenoid levels but only among controls (Figure 3C). Here, we may deduce that the exposure to LPS forced the birds with high liver carotenoid concentrations to sustain high body temperatures, flattening the regression slope (Figure 3C). However, this does not explain the lack of significant interaction in other tissues.

Among birds where the spleen carotenoids could be quantified (i.e., those with heavier spleens and paler bills), LPS-treated individuals had a higher carotenoid detection rate and a trend toward significantly higher spleen lutein levels. Despite the low sample size, both results suggest enhanced carotenoid investment in the spleen during immune activation. This supports the presence of a carotenoid allocation trade-off between immune-competence and ketocarotenoid-based signaling if we consider the LPS-induced color decline in the full sample. However, that color decrease was undetected in the reduced subsample, perhaps due to its lower size. Further evidence for strategic carotenoid allocation comes from the significant positive correlations between recent food intake and carotenoid levels in plasma and fat of LPS-treated birds, but not controls. These immune-challenged individuals may have increased carotenoid absorption and storage from food, but not the carotenoid allocation to the signal (LPS-treated birds showed paler signals).

The positive relationship between final bill redness and spleen carotenoid levels further supports the reliability of bill color as an indicator of condition. Likewise, birds with measurable carotenoid concentrations in subcutaneous fat displayed redder bills at the end of the experiment. Carotenoid storage in fat is unlikely to serve as a reservoir for re-circulation. For example, in the Greylag geese (*Anser anser*), carotenoids stored in subcutaneous fat do not seem to be released back into circulation (Negro et al., 2001). Instead, mammalian studies suggest that carotenoids are relevant in lipid reserves, functioning as antioxidants supporting lipid catabolism pathways (Bonet et al., 2015; Mounien & Tourniare, 2019). Consequently, birds with redder bills may not only indicate their ability to invest carotenoids in their immune system but also their capacity to maintain efficient lipid metabolism.

In summary, in male zebra finches, the red keto-carotenoid-based bill signal and several parameters related to its production mechanism (blood, liver and fat carotenoid levels) were associated with food intake. The link to quality is mostly related to final bill redness, suggesting that the signal informs about the recent physiological condition (see also Gautier et al., 2008, Ardia et al., 2010, McGraw et al., 2011, George et al., 2017). The reddest birds at the end of the study were also those with lower body temperatures and detectable subcutaneous fat carotenoids. Taken together, our findings support the view that plastic resource allocation trade-offs underlie the reliability of red bill coloration in male zebra finches, suggesting potential for strategic adjustment and even cheating strategies. Therefore, they do not seem to support a tight association of bill redness to individual quality, which should be expected for red keto-carotenoid ornaments more generally if they are rigidly constrained and unfalsifiable index signals (e.g. Weaver et al., 2017; Koch & Hill, 2018; Cantarero et al., 2020).

## Ethics note

This experiment was approved by the animal ethics government office (i.e., Consejería de Agricultura, Junta de Castilla La Mancha; approval ref. JCCM 27-2021). All the authors consent to participate.

## Acknowledgements

We thank María de los Santos Fernandez and Marta Garrido for their help in laboratory analyses.

## Authors contribution

AC and CAA designed and performed the experiment. RA and PC performed chromatography analyses and datasets on carotenoids and vitamin levels. CAA obtained funds for the project. CAA performed the stats and wrote the first version of the text. MC and PA contributed to the experimental design. AC, RM, PC and MC revised the drafts of the text.

