## Supplementary Material for "Sexual signal reliability in male zebra finches: food intake explains the impact of immune activation on carotenoid-based coloration"

***(Cantarero et al.)***

**METHODS**

*Experimental design complementary information*

Male adult zebra finches were acquired from seven suppliers placed in the Community of Madrid (Spain). The supplier identity was randomly distributed between the two treatments described below (*χ*^2^ = 5.41, df = 6, p = 0.490). Birds were individually caged under standardized conditions (temperature: mean ± range: 22° ± 1°C; daily light cycle: 16L:8D). A commercial mix of seeds (KIKI exotic birds) was used to feed the birds throughout the experiment. The mix was composed of paniset, yellow, white, red and black millets, canary seed and linseed. The composition provided by the company states: crude protein: 12.8%, Humidity 9.2%, Crude fat 6.7%, Crude fibre 9.3%, Ash 4.2%, Calcium 0.23%, Phosphorus 0.47%, Sodium 218 ppm, Vitamin A 8500 IU/kg, Vitamin D3 1500 IU/kg, Vitamin C 20 mg/kg, Vitamin E 25 IU/kg. We additionally measured its composition in terms of carotenoids, retinoids and tocopherols. This was made by HPLC, as described in the main text. Six samples of that mix were taken (mean ± SD: 5.04 ± 0.01 g). The samples were homogenized and saponified by means of KOH to release any carotenoid linked to fatty acids. The grain mix was composed of (mean values in nmol/g): α-tocopherol: 65.93, δ-tocopherol: 10.49, γ-tocopherol: 11.02, lutein: 49.90, zeaxanthin: 1.51, and three lutein derivatives: 0.642, 1.40 and 0.96. Total carotenoid content: 76.87 nmol/g.

All the birds were assigned to the treatments five days after blood sampling. No significant differences in body mass, wing length or tarsus size between the two treatments were found on the day of the start of the experiment (i.e., first injection day, see below; all Student’s *t* *p*-values > 0.45). Size-corrected body mass (usually termed as “body condition”) neither showed a treatment effect at that time (ANCOVA; treatment: *F*_1,66_ = 0.01, *p* = 0.944; wing and tarsus lengths as covariates with *p-*values < 0.01). Similarly, the bill redness (i.e. reversed hue; see below description) did not differ either (ANCOVA; treatment: *F*_1,66_ = 0.002, *p* = 0.963; bill brightness and selected bill’s area covariates with *p-*values < 0.01: estimates: -8.27 ± 0.97 and 1.58 ± 0.50, respectively). The selected bill’s area and brightness did not differ (ANCOVA's *p* > 0.47). The food intake before the experiment (see above) did not differ either (treatment: *F*_1,67_ = 0.10, *p* = 0.758; first body mass measurement: *F*_1,67_ = 16.15, *p* < 0.001, estimate ± SE: 0.26 ± 0.07). The treatment was alternatively distributed among the aviary cages, and the birds from each treatment were alternated at the time of sampling to avoid biases related to day-time cycles (e.g., body temperature; Sköld-Chiriac et al., 2015; circulating carotenoids; Horak et al., 2004; but see Pérez-Rodríguez et al., 2007).

*Experimental chronogram*

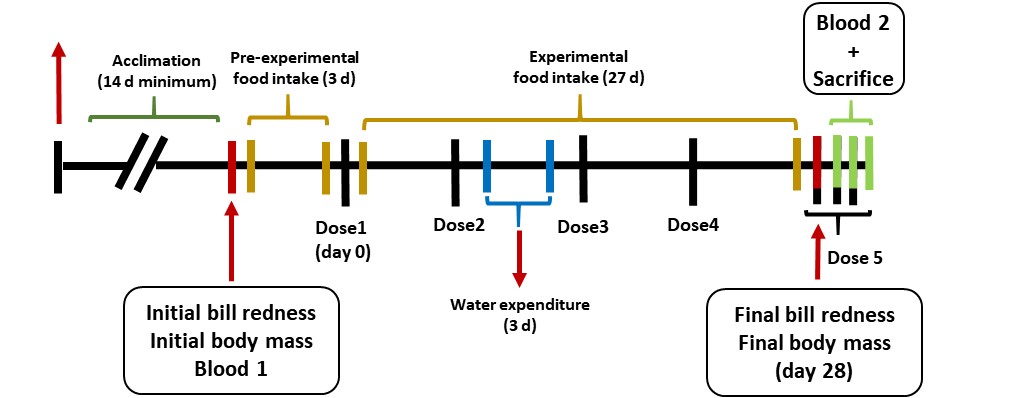

Figure S1. Experimental chronogram. Birds were housed in individual cages upon arriving from providers. After an acclimation period, a first measure of bill colour, morphometries and body mass was taken, and a blood sample was extracted. Two days later, the feeders were replenished and weighed. This was repeated every three days throughout the study. The difference in weight between the first two feeder measures was used to calculate individual pre-experimental food intake. Subsequent measures were used to determine the total experimental food intake (see also the main text). Water dispensers were weighed twice within a three-day lapse to estimate individual water expenditure during the experiment. Birds received the first injection (day 0 of the experiment) one day after the second feeder weight measurement, with injections repeated weekly. Due to time constraints for euthanasia and tissue sampling, birds were divided into three sampling blocks at the end of the experiment. Final measures of bill redness and body mass were taken for all birds the day before the first sacrifice.

*Colour measurements*

The same standard grey reference and scale (ColorChecker Classic target; X-Rite, Michigan) was placed next to the bird's head for each photo. The focus and diaphragm of the camera were manually fixed to avoid the interference of automatic functions. We have previously shown that these picture-based measurements are highly correlated with the redness measurement (i.e., red hue) obtained from portable spectrophotometers (Alonso-Alvarez & Galván, 2011; Mougeot et al., 2007). SpotEgg software, however, allows the user to manually draw any region and provide information about its colouration, shape or other features. As opposed to portable spectrophotometers that analyze the colouration of reduced spots (usually 1-2 mm), the measure of a large area makes this tool useful for evolutionary biologists aiming to capture most of the variability among individuals (Gómez & Liñan-Cembrano et al., 2017). Accordingly, the average of red, green, and blue (RGB) components of the lateral bill surface (upper and lower mandibles) were calculated for each animal. We used the lateral side of the bill due to the low variance of colouration at the top. We then determined hue values using the Foley & van Dam algorithm (1982) algorithm. Since a low hue measure means a redder colour, the hue value was reversed. This reversion was made by multiplying hue values by -1 and adding the minimum value to attain positive data. The new value was defined as “redness”.

*Carotenoid extraction*

Liver (mean 50 mg), spleen (mean 33 mg) and subcutaneous fat (mean 90 mg) samples were vortexed for 1 min in a cryotube (2 mL) with stainless steel balls with 1 mL of a mixture of methanol : water (1 : 1). An internal standard (IS) solution (50 µl) containing retinyl acetate (50 uM) and tocopheryl acetate (30 mM) (both from Sigma-Aldrich) was also added to each sample. Then, 500 µL of hexane was added, and it was vortexed again for 5 min and centrifuged at 14000 rcf for 5 minutes. The supernatant (≈ 480 µl) was transferred to a glass tube, kept on ice by avoiding light exposure, and evaporated to dryness under a nitrogen stream. The dry extract was redissolved in 200 µL of the initial chromatographic phase formed by methanol : methyl tert-butyl ether : water (80 : 13 : 7). In case of the beak, the sample (mean 8 mg) was homogenized with 5 mL of methanol : water (1 : 1) in the steel capsule of a mixer mill (Retsch MM400) with a steel ball and with the same amount of IS as the other tissues. The obtained extract was transferred to a glass tube, and 2 mL of hexane was added, gently shaken for 1 min, centrifuged at 2000 rcf for 5 minutes, and the supernatant was processed as for the rest of the tissues. Plasma samples (0.05 ml) were extracted following García-de Blas et al. (2013) with 150 µl of ethanol and 50 µl of water, and it was processed as the rest of the samples, including the same amount of IS.

**COMPLEMENTARY RESULTS**

The total amount of food consumed by each bird positively correlated with the initial body mass measured when the recording of food consumption started (Figure S2).

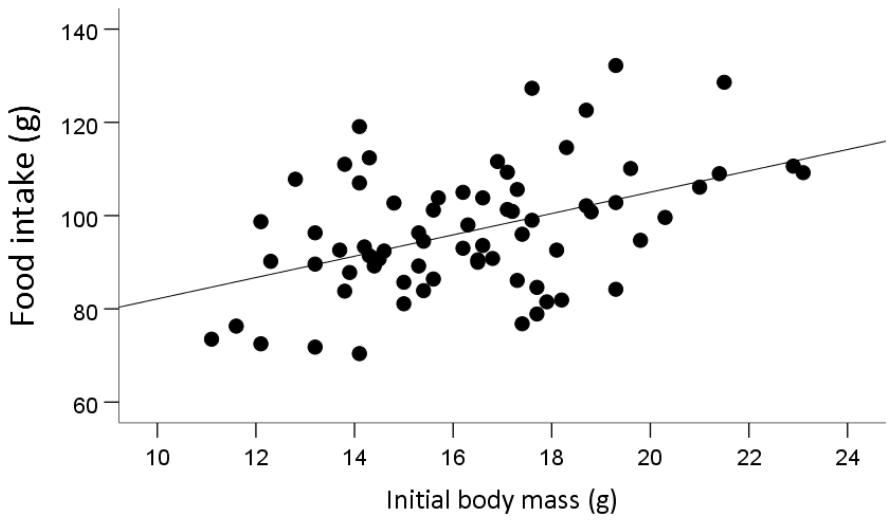

Figure S2. Heavier birds consumed more food (grain) during the experiment than lighter individuals (*R*^2^ = 0.20, *p* < 0.001, slope ± SE: 2.29 ± 0.55).

The bill total carotenoids were positively correlated to bill redness (reversed hue) at the end of the experiment (Figure S3).

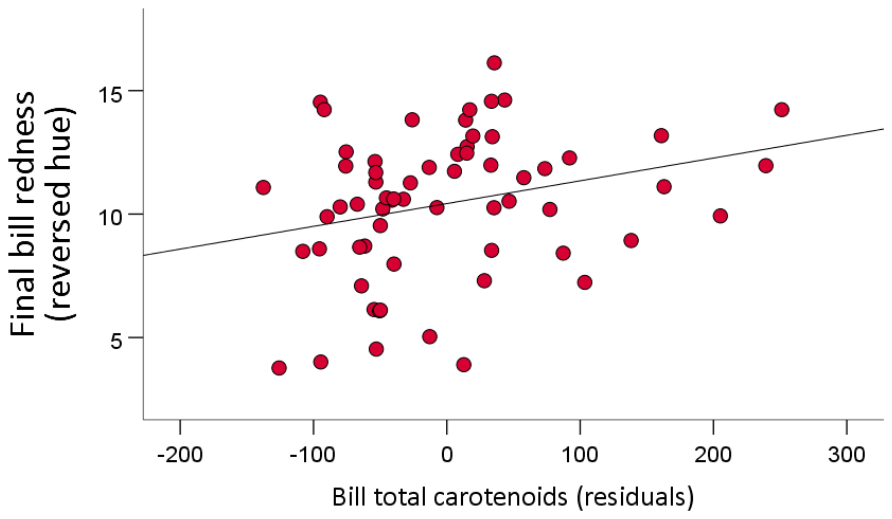

Figure S3. Relationship between bill redness (reversed hue) at the end of the experiment (sacrifice day) and bill total carotenoid levels (residuals from the mixed model; see main text).

The liver's total carotenoid concentration positively correlated with food intake (Figure S4).

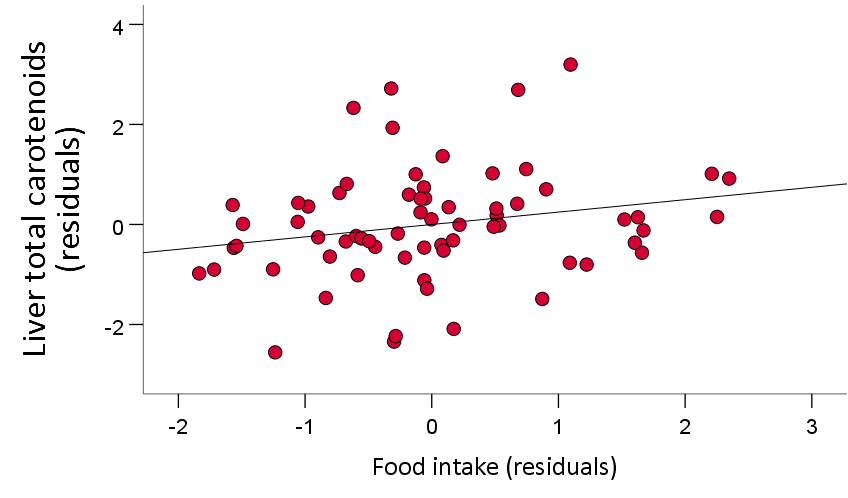

Figure S4. Liver total carotenoids correlated to food intake. Both variables are residuals from models described in the main text.

Regarding spleen carotenoids, echinenone and canthaxanthin peaks were also identified in two birds only (one control and one LPS bird, the same birds in both pigments). With this scarce data, we decided to exclude them from the statistical analyses.

The relationship between initial bill redness and spleen mass measured approximately one month later was also significantly positive in the reduced subsample of birds where the spleen carotenoids could be quantified (Figure S5).

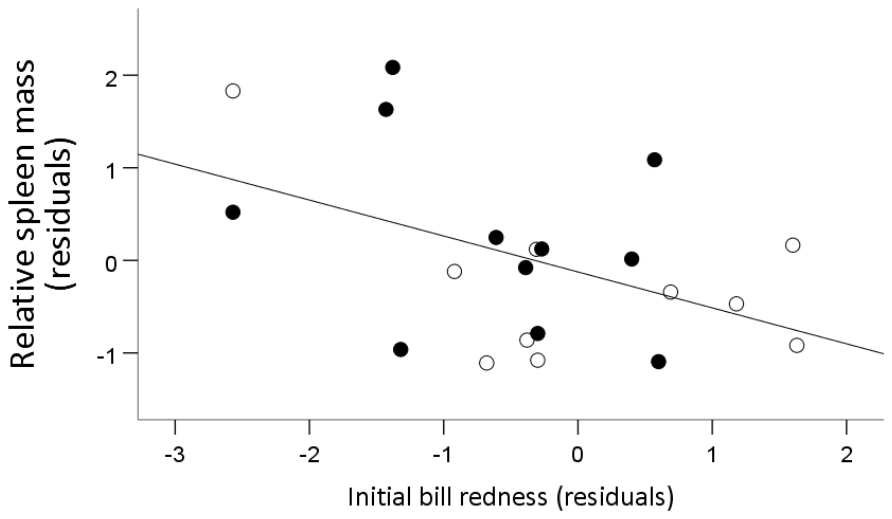

Figure S5. Relationship between relative spleen mass (corrected for body mass at sacrifice) and initial bill redness (controlled for initial bill brightness and area) in the subsample of birds that provided enough spleen tissue for carotenoid quantification (*r* = -0.46, *P* = 0.035; *n* = 21). Empty dots for controls and filled dots for LPS-treated birds.

*Alternative results with raw body temperatures*

We repeated the analyses of body temperature excluding those nine birds that provided low values due to the measurement method (see Methods in the main text). In this subsample (*n* = 61), the treatment showed a trend to a significant effect with LPS-treated birds showing higher temperatures than controls (*t* = 1.81; df = 59, *p* = 0.075, n = 61; *d* = 0.46; Figure S6).

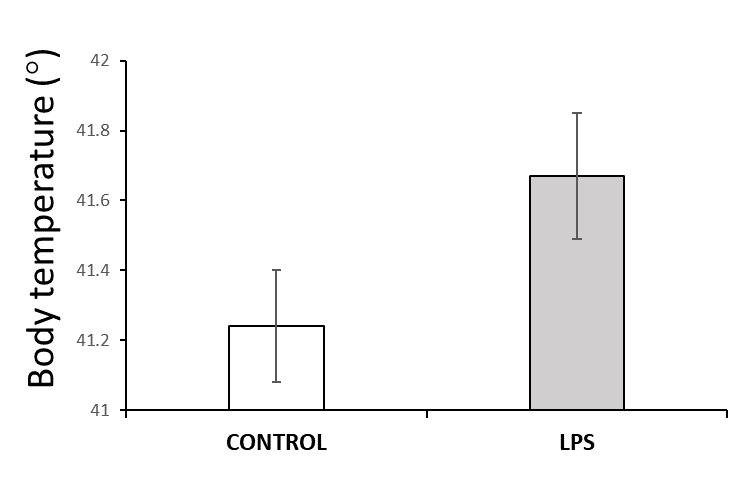

Figure S6. Body temperature (raw values) depending on treatment (*n* = 61; means ± SE).

The relative spleen mass was uncorrelated with the body temperatures (*r* = -0.06, *p* = 0.638, *n* = 61). We also tested the body temperature as a covariate in the Table 1 bill redness model, reporting a negative correlation (*F*_1.55_ = 8.86, *p* = 0.004; slope ± SE: -0.925 ± 0.311). Redder bill bills showed lower temperatures. That link remained when removing the initial bill redness, i.e. testing the colour at sacrifice independent of initial variability (*F*_1.56_ = 5.08, *p* = 0.015; slope ± SE: -0.869 ± 0.345). Again, the redder the bill, the lower the temperature (*r* = 0.34, *p* = 0.007, *n* = 61; Figure S7A). Body temperature did not interact with treatment on bill redness (both *p-*values > 0.70).

In the model testing liver total carotenoids, the body temperature covariate also showed an interaction with treatment close to significance (*F*_1.53.6_ = 1.59, *p* = 0.213; interaction: covariate: *F*_1.47_ = 3.56, *p* = 0.052). There was no relationship in LPS-treated individuals (*F*_1.21.4_ = 0.36, *p* = 0.554; slope ± SE: 0.132 ± 0.219, *n* = 26), but a negative correlation close to significance in controls (*F*_1.29.2_ = 4.12, *p* = 0.052; slope ± SE: -0.523 ± 0.258; *n* = 32; Figure S7B).

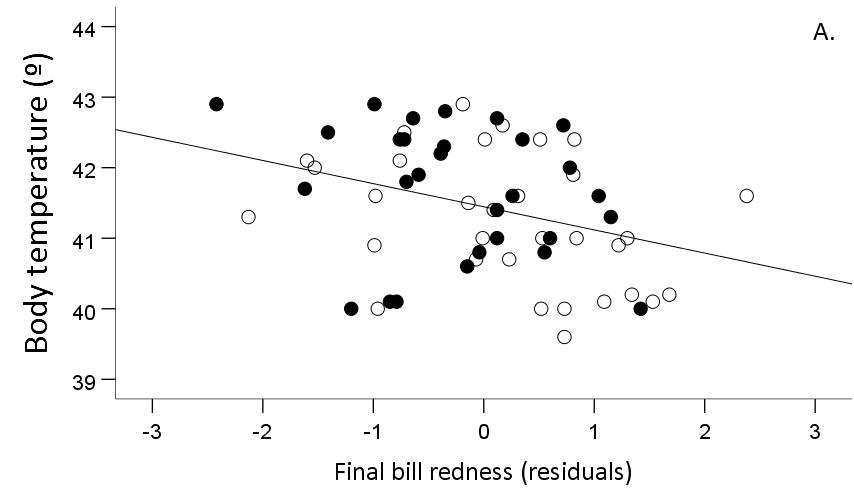

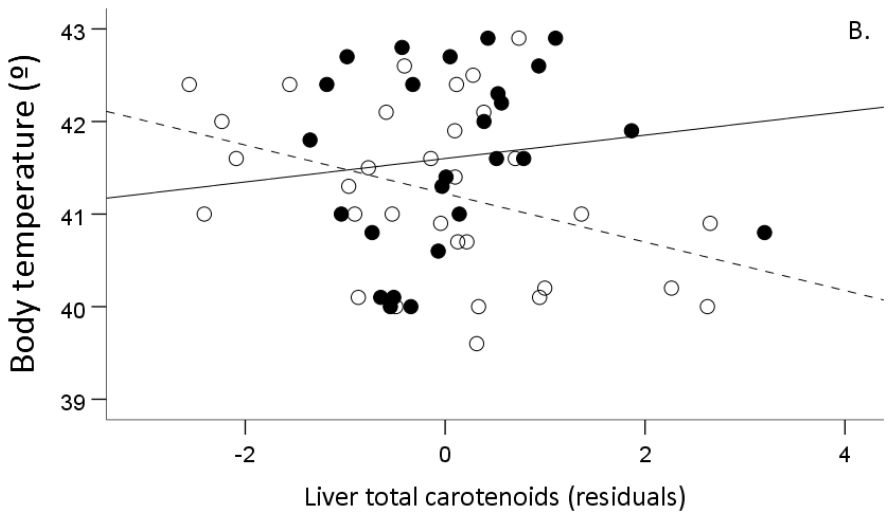

Figure S7. Relationship between body temperature and final bill redness (A) and liver total carotenoids (B).
