## Supplementary material for "Sexual signal reliability in male zebra finches: food intake explains the impact of immune activation on carotenoid-based coloration": Table S1

**Table S1.** Carotenoid and vitamin composition of different male zebra finch tissues. The parameters are reported in elution order.

| **Bill (nmol/g)** | **N** | **Minimum** | **Maximum** | **Mean** | **SD** |
| --- | --- | --- | --- | --- | --- |
| Retinol | 63 | 0.00 | 8.60 | 0.48 | 1.46 |
| γ-tocopherol | 63 | 0.03 | 506.95 | 204.54 | 97.65 |
| α-tocopherol | 63 | 2.46 | 1373.63 | 528.45 | 250.79 |
| Zeaxanthin | 63 | 0.04 | 28.41 | 10.64 | 5.74 |
| α-doradexanthin | 63 | 0.31 | 24.68 | 9.45 | 4.77 |
| Astaxanthin | 63 | 2.68 | 303.35 | 116.52 | 63.07 |
| Unidentified carotenoid 1 | 63 | 0.04 | 31.31 | 9.65 | 5.67 |
| Adonirubin | 63 | 0.28 | 27.55 | 11.07 | 5.50 |
| Unidentified carotenoid 2 | 63 | 0.09 | 58.75 | 24.67 | 11.61 |
| **Blood 1 plasma (nmol/mL)** | **N** | **Minimum** | **Maximum** | **Mean** | **SD** |
| Retinol | 67 | 11.04 | 58.57 | 31.33 | 10.94 |
| Lutein | 67 | 4.04 | 177.82 | 48.67 | 33.32 |
| Lutein derivative 1 | 67 | 0.00 | 22.99 | 5.20 | 4.74 |
| Lutein derivative 2 | 67 | 0.00 | 11.42 | 2.51 | 2.28 |
| Zeaxanthin | 67 | 0.00 | 8.80 | 2.18 | 1.85 |
| Lutein derivative 3 | 67 | 0.00 | 138.50 | 34.82 | 25.86 |
| Lutein derivative 4 | 67 | 0.00 | 26.43 | 6.76 | 5.37 |
| γ-tocopherol | 67 | 0.00 | 145.84 | 12.73 | 28.16 |
| α-tocopherol | 67 | 1.89 | 232.16 | 22.30 | 47.57 |
| **Blood 2 plasma (nmol/mL)** | **N** | **Minimum** | **Maximum** | **Mean** | **SD** |
| Retinol | 70 | 9.00 | 69.80 | 32.70 | 12.31 |
| Lutein | 70 | 4.08 | 192.39 | 49.05 | 33.41 |
| Lutein derivative 1 | 70 | 0.00 | 22.87 | 5.35 | 4.45 |
| Lutein derivative 2 | 70 | 0.00 | 12.63 | 2.57 | 2.33 |
| Zeaxanthin | 70 | 0.00 | 10.09 | 2.34 | 2.02 |
| Lutein derivative 3 | 70 | 0.00 | 136.90 | 36.91 | 26.43 |
| Lutein derivative 4 | 70 | 0.00 | 26.66 | 6.68 | 5.19 |
| Echinenone | 70 | 0.00 | 0.64 | 0.02 | 0.09 |
| γ-tocopherol | 70 | 0.00 | 144.65 | 12.60 | 27.56 |
| α-tocopherol | 70 | 1.77 | 260.59 | 24.19 | 53.85 |
| **Liver (nmol/g)** | **N** | **Minimum** | **Maximum** | **Mean** | **SD** |
| Retinol | 67 | 7.52 | 33.95 | 15.02 | 5.77 |
| γ-tocopherol | 67 | 13.25 | 29.81 | 19.93 | 4.03 |
| α-tocopherol | 67 | 18.41 | 74.02 | 40.79 | 14.21 |
| Lutein derivative 1 | 67 | 2.17 | 2.39 | 2.27 | 0.05 |
| Lutein | 67 | 2.23 | 3.03 | 2.49 | 0.18 |
| Zeaxanthin | 67 | 2.25 | 3.49 | 2.63 | 0.26 |
| Lutein derivative 2 | 67 | 0.00 | 3.40 | 2.39 | 0.64 |
| β-cryptoxanthin | 67 | 2.58 | 4.59 | 3.12 | 0.42 |
| Lutein derivative 3 | 67 | 0.00 | 2.39 | 2.24 | 0.28 |
| Retinoid 1 | 67 | 7.79 | 95.73 | 31.53 | 20.30 |
| Retinoid 2 | 67 | 9.31 | 209.51 | 42.88 | 34.09 |
| Retinoid 3 | 67 | 55.84 | 1306.12 | 450.71 | 302.84 |
| Retinoid 4 | 67 | 4.00 | 207.79 | 58.24 | 47.61 |
| **Fat (nmol/g)** | **N** | **Minimum** | **Maximum** | **Mean** | **SD** |
| Retinol | 70 | 0.00 | 57.38 | 9.29 | 9.82 |
| g- tocopherol | 70 | 0.00 | 126.23 | 28.77 | 20.72 |
| α-tocopherol | 70 | 0.00 | 66.87 | 17.96 | 14.33 |
| Lutein | 70 | 0.00 | 7.70 | 1.46 | 1.67 |
| Zeaxanthin | 70 | 0.00 | 17.37 | 2.43 | 3.21 |
| Lutein derivative 3 | 70 | 0.00 | 8.72 | 0.85 | 1.86 |
| **Spleen (nmol/g)** | **N** | **Minimum** | **Maximum** | **Mean** | **SD** |
| Retinol | 21 | 0.45 | 37.59 | 7.27 | 10.20 |
| Lutein derivative 3 | 21 | 0.00 | 8.37 | 1.84 | 2.62 |
| Lutein derivative 1 | 21 | 0.00 | 8.37 | 1.65 | 2.60 |
| Lutein | 21 | 0.00 | 4.08 | 1.65 | 1.16 |
| Zeaxanthin | 21 | 0.00 | 9.69 | 1.59 | 2.38 |
| Cantaxanthin | 21 | 0.00 | 0.27 | 0.01 | 0.06 |
| Lutein derivative 2 | 21 | 0.00 | 11.34 | 3.69 | 3.61 |
| Retinoid 1 | 21 | 0.00 | 110.56 | 17.12 | 33.46 |
| Retinoid 2 | 21 | 0.00 | 179.53 | 27.06 | 52.23 |
| β-cryptoxanthin | 21 | 0.00 | 6.55 | 0.88 | 1.77 |
| Echinenone | 21 | 0.00 | 0.71 | 0.07 | 0.21 |
| Retinoid 3 | 21 | 0.00 | 2587.56 | 313.29 | 655.53 |
| Retinoid 4 | 21 | 0.00 | 288.15 | 40.25 | 80.34 |
| γ-tocopherol | 21 | 0.00 | 54.20 | 23.05 | 14.18 |
| α-tocopherol | 21 | 0.00 | 54.29 | 24.19 | 15.18 |
